# Anemia Diagnosis on a Simple Paper-based Assay

**DOI:** 10.1101/439224

**Authors:** Sujay K Biswas, Soumya Bandyopadhyay, Shantimoy Kar, Nirmal K Som, Suman Chakraborty

## Abstract

In developing countries, the maternal and neonatal mortality rate is often affected by prenatal period anemia, a preventable and ubiquitous impairment attributed due to low hemoglobin (Hgb) concentration. We report the development of a simple, frugal (~ 0.02 $ per test), rapid and high fidelity paper-based colorimetric microfluidic device for point-of-care (POC) detection of anemia. We validate our findings with 32 blood samples collected from different patients covering a wide spectrum of anemia and subsequently, compare with standard pathological results measured using a hematology analyzer. POC based Hgb estimates are correlated with the pathological gold standard estimates of Hgb levels (r = 0.909), and the POC test method yielded similar sensitivity and specificity for detecting mild anemia (n = 8) (<11 g/dl) (sensitivity: 87.5%, specificity: 100 %) and for severe anemia (n = 3) (<7 g/dl) (sensitivity: 100 %, specificity: 100 %). The estimated Hgb levels are, within 1.5 g/dl from the pathological estimate, for 91 % of the blood samples. Results demonstrate the elevated efficacy and viability of this POC colorimetric diagnostic test, in comparison to the state-of-the-art complex and expensive diagnostic tests for anemia detection.

## Introduction

Anemia, a type of blood disorder, is perceived as one of the most prevalent diseases which has severely affected one-third of the world’s population (2.36 billion people) (1, 2). The widespread causes behind anemia are fundamental hematologic ailments (e.g. sickle-cell disease, thalassemia), iron deficiency, dietary deficiencies, and chronic conditions (e.g. HIV, kidney disease, cirrhosis, inflammatory immune disorders) (3–9). Anemic patients not only experience fatigue, dizziness, and severe dehydration, but also suffer from critical complications like angina, congestive heart failure, heart diseases, and stroke(10). Anemia can trigger abated work capacity, retarded cognitive and physical growth, undesirable preterm delivery, under-weight babies and thus augmented mortality risk. According to WHO, the lower threshold of hemoglobin for anemia to be 13 g/dl for men, 12 g/dl for non-pregnant women, and 11.5 g/dl for children ages 5.0-11.9 years, 11 g/dl for pregnant women and young children ages 0.5 - 4.99 years (11). Anemia is diagnosed by estimating the hemoglobin level in blood, which, in turn, is dictated by the count of healthy RBCs. Early diagnosis may reduce the risk of further health complications by initiating standard treatment with necessary vitamin and dietary iron supplements. However, in the developing world, still, there is a significant dearth of rudimentary medical facilities, catering to the entire population. The paucity of clinical amenities, in conjunction with the lack of awareness, has further aggravated the current situation. Consequently, this problem attains utmost importance and entails attention from the broad scientific community.

Pathology labs (12, 13) ubiquitously employ cyanmethemoglobin as a standard and Drabkin’s solution as the reagent for spectrophotometric quantification of hemoglobin levels. Kaiho and Muzino (14) expounded sensitive and rapid colorimetric assays based on 2,7- diaminofluorene. Toh et al. (15) used glassy carbon electrodes coated with Nafion layers for electrochemical detection of hemoglobin by invoking the principle of cyclic voltammetry. Chitnis et al. (16) fabricated a microfluidic substrate by etching hydrophilic patterns on a hydrophobic coated paper substrate using laser treatment by CO_2_ laser and subsequently detected hemoglobin by adopting a luminol-based chemical pathway. Barati et al. (17) employed carbon dots as a fluorescent probe for conducting spectrofluorometric hemoglobin detection from DI water diluted blood. A multi-step procedure of scanner based hemoglobin detection method by deploying dried stain of lysed blood on chromatography paper has been reported by Yang et al (10). Taparia et al. used PDMS microchannel to detect the level of hemoglobin optically by using a CMOS sensor from diluted blood with calcium-free Tyrode buffer (18). It is undeniable that a significant progress has been made towards the development of a better methodology for point-of-care hemoglobin detection. However, none of these techniques are cost-effective and suitable for use in a resource-limited setting, still necessitate highly expensive fabrication procedure, complex detection methodology, a large volume of sample and reagent, and trained personnel. Hence, there is a need for an affordable, simple, rapid and high fidelity diagnostic platform which can be employed as a counteractive measure for the paucity of conventional pathological diagnostic platforms.

In this present work, we have demonstrated a frugal, user-friendly, and paper-based point-of-care microfluidic device for colorimetric detection of anemia. Our paper is the first report where we can reproduce gold standard results from the pathological laboratory by improvising and innovating adapted simple chemical protocols on an ultra-low-cost microfluidic chip (essentially, a piece of paper), taking cues from the fundamental chemistry governing the underlying colorimetric detection. In sharp contrast to other reported point of care devices for hemoglobin level detection, this device turns out to be of significant low cost, deployable with minimally trained personnel in extreme point of care settings, and capable of reproducing gold standard results from a quantitative perspective. In essence, thus, our innovative methodology not only adopted an inexpensive simple fabrication procedure with prevalent paper substrate but also enables faster and more precise quantitative detection of hemoglobin levels using minimal quantities of blood, by adopting straightforward chemical assays and uncomplicated detection algorithms, compared to existing approaches (10, 19–21). For instance, Bond et al. (22) adopted an integrated approach in quantifying hemoglobin concentrations in blood using spectrophotometer validated chip fabricated from chromatography paper. However, the delineated approach entails the involvement of expensive and cumbersome appliances and excessive quantities of blood, thereby increasing the complexity of the detection protocol. We have not only been able to surpass the accuracy of the ubiquitously used diagnostic platforms, e.g. WHO Hemoglobin Color Scale, with our proposed methodology, but also have significantly minimized the associated expenses. Furthermore, the developed frugal microfluidic platform can be easily adapted for mass production in the industry due to the straightforward manufacturing protocol, thus can be up-scaled efficiently.

## Results and Discussion

The colorimetric assay involves a Hgb catalyzed redox reaction between 3,3′-Dimethyl-[1,1′- biphenyl]-4,4′-diamine (o-tolidine) and hydrogen peroxide yielding greenish-blue colored oxidized o-tolidine products (23). This reaction is mechanistically similar to the well-established and commercially adopted chemical assay which has been utilized in the spectrophotometric detection of Hgb in plasma, criminal inspection, and in portable POC photometers for anemia diagnosis (19, 24, 25). We have readjusted and altered the reaction parameters (please see Supplementary Information) to obtain quantifiable colorimetric signals, corresponding to the Hgb levels spanning the entire physiological range; thereby recasting this assay into a rapid, reliable and point-of-care colorimetric detection of Hgb and subsequent diagnosis of anemia without any ancillary equipment.

We have explicated the kinetics of the involved chemical reaction while calibrating our paper-based microfluidic device. Reaction kinetics (Figure 2B), is depicted by the temporal variation of the normalized intensity, obtained by: 
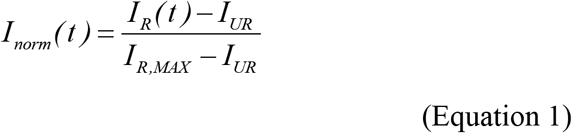

**Figure 1.**
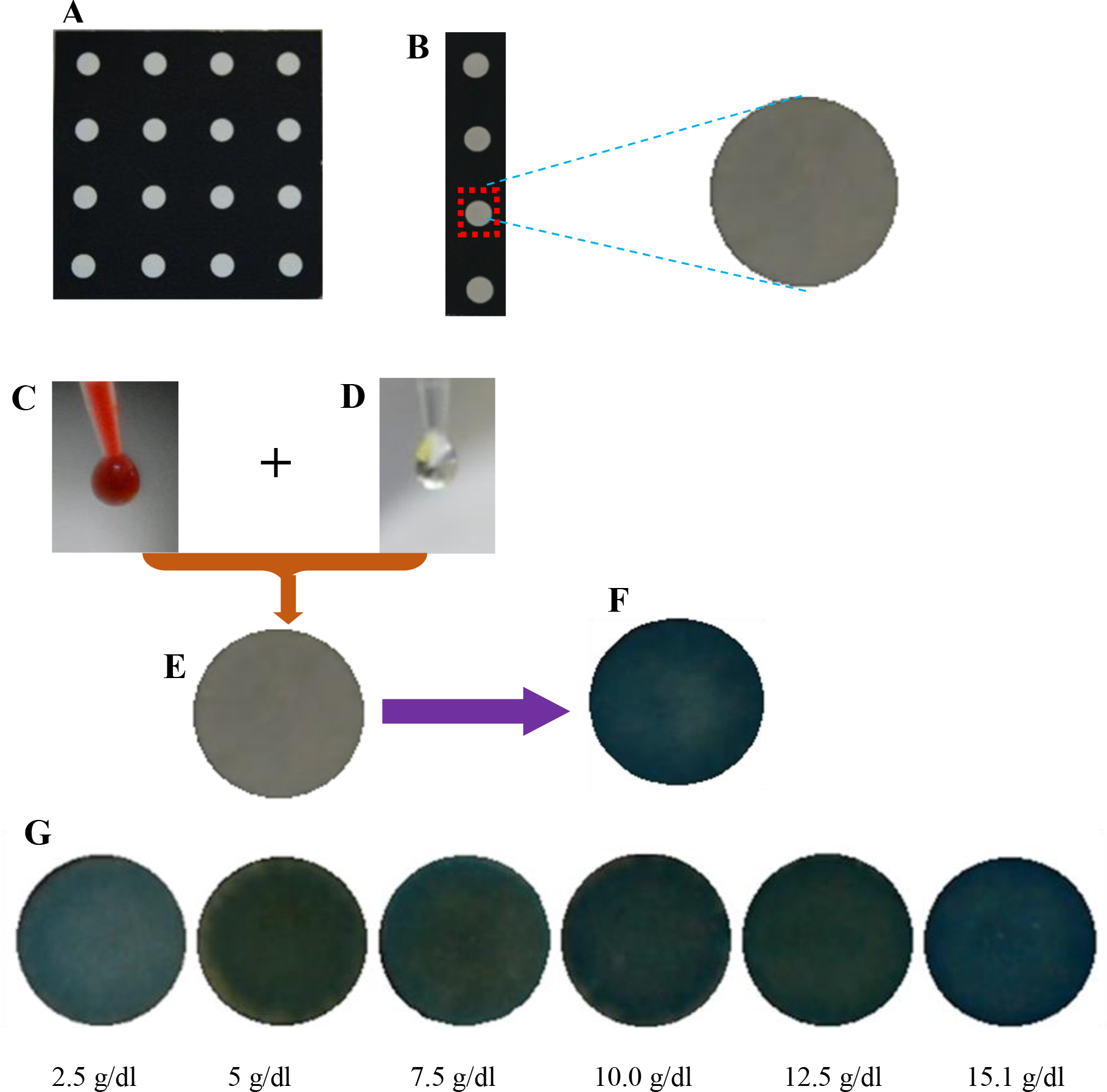
The POC color-based anemia test. (**A**) The microfluidic POC chip comprises 16 circular detection sites of diameter 0.7 cm each, separated by a hydrophobic barrier, in a square array (8 cm × 8 cm). (**B-F**) Steps for using the POC color-based anemia test. **(B)**Depicts the magnified view of a reagent immobilized reaction site. (**C**) Blood, mixed with Drabkin’s solution (**D**) added to the detection sites **(E). (F)**Wait until complete coloration and interpret developed color of the detection site. (**G**) Ranging from lighter shade of bluish green to deeper shade, resultant solution colors map with different Hgb levels, as outlined.

**Figure 2.**
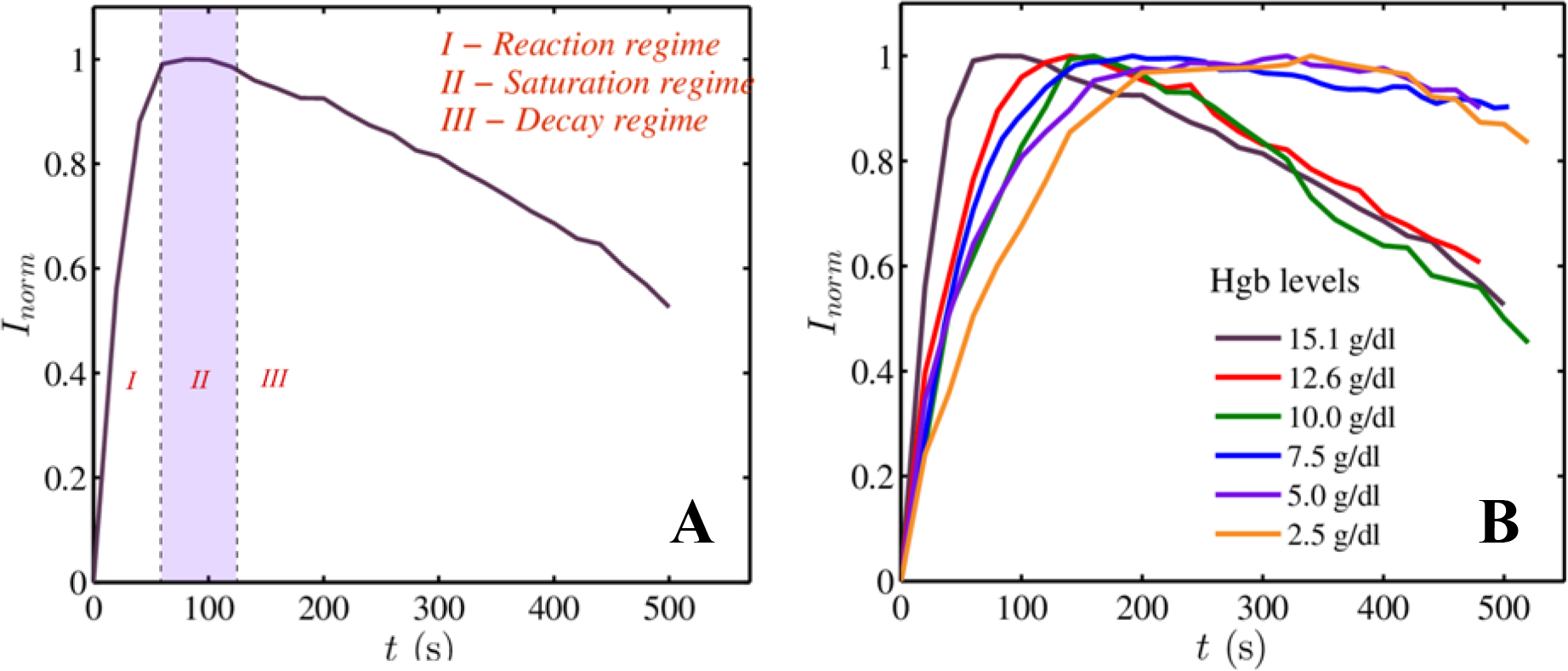
Quantification of the reaction kinetics of POC device for color-based anemia test. **(A)**Plot depicting the variation of normalized intensity (*I*_*norm*_) with time (*t*). Different regimes of colorimetric reaction are outlined. Normalized intensity maxima lie in the regime *II.* **(B)**Curves depicting reaction kinetics of resultant cyanmethemoglobin solution for blood samples with Hgb levels of 2.5 g/dl, 5.0 g/dl, 7.5 g/dl, 10.0 g/dl, 12.6 g/dl, and 15.1 g/dl.

The temporal variables *I*_*norm*_(*t*), *I*_*R*_(*t*), *I*_*R,MAX*_ and *I*_*UR*_ denote the normalized intensity, the intensity of the reacted pad, the average intensity of saturated color and intensity of the unreacted pad respectively. The reaction kinetics, for a particular concentration, in general, comprises three distinct phases (Figure 2A). The reaction regime (*I*) signify chemical reaction till its completion. The shaded regime (*II*) is the temporally stable saturation region; the temporal average of the intensity in this region (*I*_*R,MAX*_) has been utilized for obtaining the calibrated intensity level (Δ*I*).

In the decaying regime (*III*), the colorimetric signal gradually fades which impacts in the decline of the intensity of the reaction pad. Construction of the calibration curve is performed by executing repeated experiments with the serial dilutions of cyanmethemoglobin, obtained by the treatment with standard Drabkin’s solution. The difference of the scaled intensities (Δ*l*) is obtained for a particular trial, corresponding to a given concentration, by invoking the previously discussed formulation. The mean and the standard deviation corresponding to a set of experimental trials is obtained after filtering out the outliers outside two standard deviations from the mean and subsequently mapped with the given concentration (linear fit yields r = 0.977) (Figure 3).

**Figure 3.**
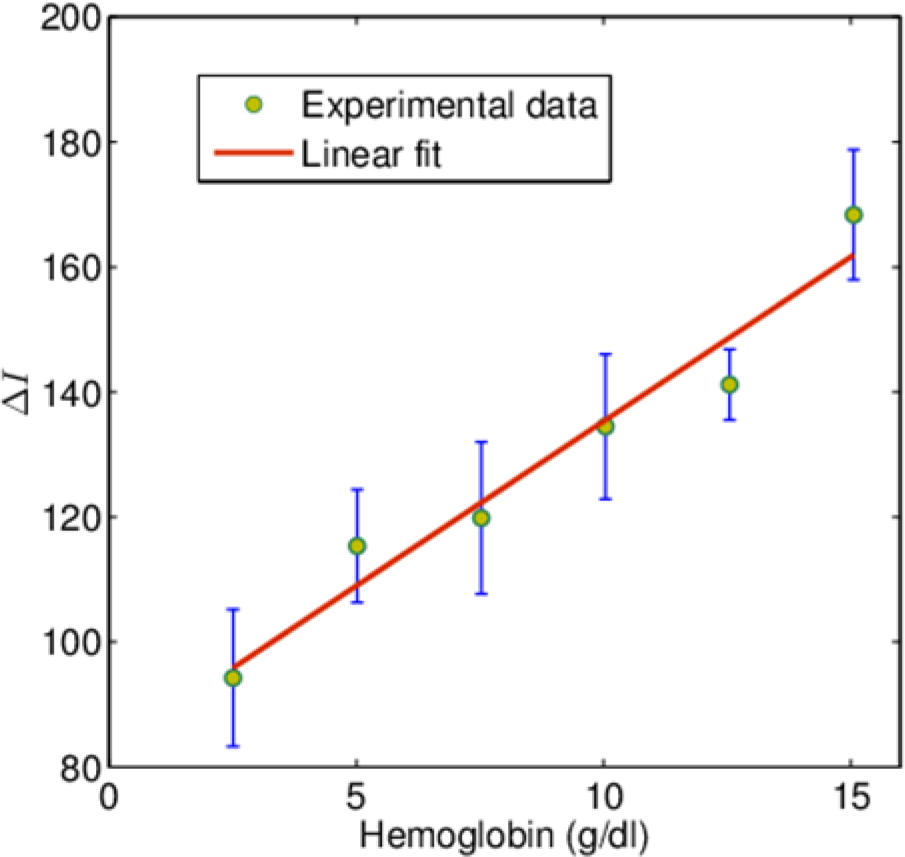
Standard calibration curve of POC device for color-based anemia test. A standard curve was constructed using serial dilutions of standard hemoglobin to create Hgb levels ranging between 2.5 g/dl and 15.1 g/dl. *R* value is 0.977. The error bars in the plot represent the standard deviation in the mean intensity computed from 20 trials corresponding to a given dilution.

Our calibrated microfluidic platform is tested with human blood samples covering the entire physiologic range of Hgb. Our device has shown a significantly high degree of repeatability, corroborated by the measured intensity of the colorimetric signals, whose coefficient of variation lies below 10 % (for 97 % samples) (Supplementary Figure 1). The concentration of hemoglobin has been subsequently estimated from the measured intensity value, by appropriately evoking the calibration curve. Our POC anemia test obtained results, which exhibit strong correlation to those of hematology analyzers (*r* = 0.909) (Figure 4), and yielded comparable sensitivity and specificity for detecting anemia (<11 g/dl) (sensitivity: 87.5%, specificity: 100%) and severe anemia (<7 g/dl) (sensitivity: 100%, specificity: 100%), equivalent to the state-of-the-art POC anemia diagnostic tests (19, 26–33). We have estimated the hemoglobin concentration, approximately within 1.5g/dl from the pathological estimate, for 91% of the blood samples. The ubiquitously used WHO Hemoglobin Color Scale compares the color of a blood spot on paper with reference standards ranging from 4 g/dl to 14 g/dl in intervals of 2 g/dl to quantify hemoglobin levels. In laboratory settings, the same color scale has displayed 95 % agreement within the limits of −3.50 g/dl to +3.11 g/dl (33). This technique is susceptible to inaccuracies: Van den Broek et al. reported that the color scale yielded quantitative measurements within 2 g/dl for only 67 % of the samples (34). Our device has not only performed better in comparison to the widely used WHO Hemoglobin Color Scale Method (33, 34) in similar conditions but also has displayed an elevated degree of efficacy compared to other low-cost devices reported in the literature(20, 22). Results reveal the potency of our microfluidic methodology to yield a greater degree of accuracy over the existing methods with curtailed cost, in similar settings.

**Figure 4.**
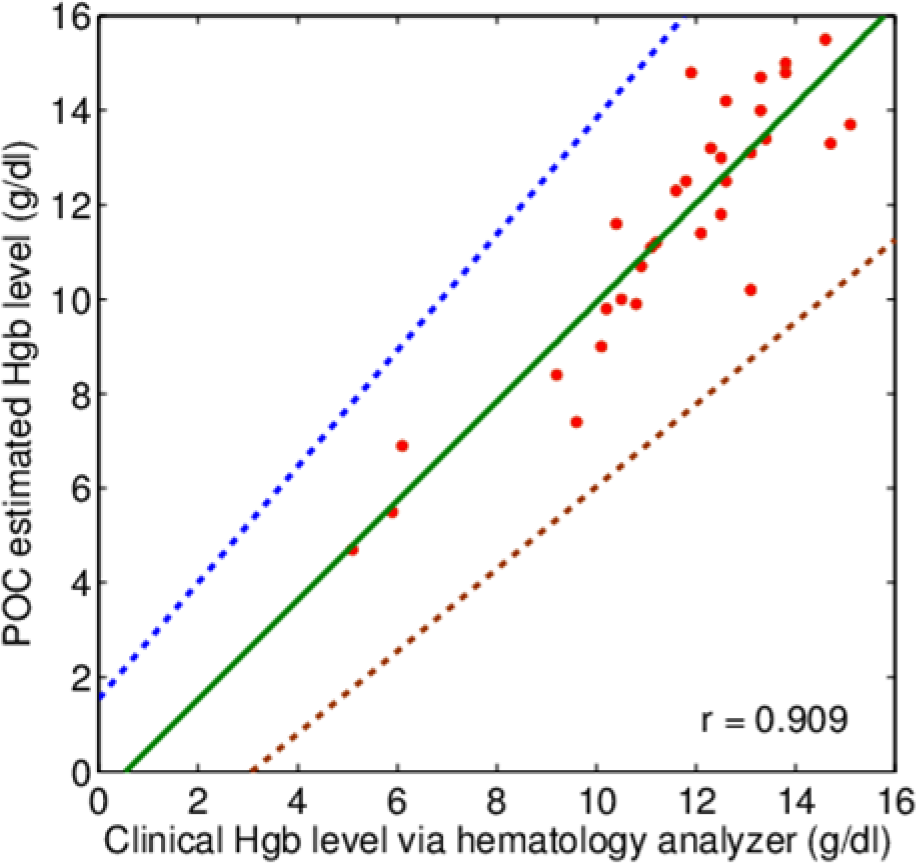
Clinical assessment of POC colorimetric anemia test results. POC Hgb levels are plotted against clinical Hgb levels determined via a hematology analyzer for all patient samples (*n* = 32), depicting a strong correlation (*r* = 0.909). Furthermore, 95% prediction limits for the individual values were computed (dashed lines).

The combined cost of Whatman 1 filter paper, extracted to 8 cm × 8 cm and combined with the requisite reagents, was estimated to have an approximately 99% reduction in expenses, relative to the HemoCue cuvettes which were priced at $0.99 - $1.43 per cuvette (as of January 2013), depending on the organization from which it is purchased (20). In comparison, the aggregate cost of an 8 cm × 8 cm strip of the Whatman Paper and the required chromogenic reagent per test contributes to $0.0123, which is a 99.4% - 99.6% depreciation in terms of price from HemoCue cuvettes. The expenses incurred in printing the pattern, followed by heating the patterned paper substrate, amount to $0.0081 per test. Hence, the aggregate cost including the material and the manufacturing cost, for the hemoglobin strip is $0.02 (Supplementary Table 1). Considering the aggregate cost, our device is more inexpensive than the existing low-cost point-of-care devices reported in the literature (20, 33, 34). We can reduce the expenses significantly by adopting a rapid prototyping manufacturing protocol. Our eco-friendly paper microfluidic chip, as opposed to the glass cuvettes used in pathological labs, does not necessitate intricate waste management strategies, and it can be conveniently incinerated in situ to discard detrimental biohazards.

In conclusion, we have demonstrated a simple, affordable, rapid and high fidelity method of detecting hemoglobin from whole blood. Our paper-based platform has displayed greater sensitivity and specificity for the entire spectrum of anemic condition, relative to the prevailing methodologies; thus could potentially bridge the lacuna in the healthcare network for the resource-limited settings.

## Methods

### Fabrication of the POC device

Microfluidic paper strips were fabricated on Whatman (Grade-1) cellulose filter paper [mean pore diameter (*2r*_*m*_): 11 μm, *r*_*m*_ stands for the mean pore radius], by ink-jet printing-based procedure, an indigenous technique developed by our group (35). Our methodology incorporates cartridge ink for creating the hydrophobic barrier surrounding the circular detection pads. The fabrication methodology is relatively simpler, frugal and suitable for large-scale manufacturing, compared to existing techniques like wax-printing (36, 37) and use of complex reagents for creating barriers (38, 39). In the adopted methodology, the prototype of the chip, to be employed, is printed on both sides of the substrate, by an ink-jet printer (HP Colour LaserJet 500). Finally, each side of the printed paper is heated to a temperature of 180ºC for 4-5 minutes. While heating the printed paper, toner particles melt, and molten ink diffuses through the porous paper matrix; thereby generating the hydrophobic barrier spanning across the paper thickness. Amidst the hydrophobic surface, hydrophilic reaction pads for colorimetric detection are situated, which will guide the flow of test fluid through the porous network of the paper matrix. The final fabricated prototype (Figure 1A) comprises 16 hydrophilic sites, which serve as the “reaction pads”.

### Color-based POC Anemia testing

We calibrated the fabricated microfluidic platform with different dilutions of cyanmethemoglobin, which were prepared by mixing varied ratios of cyanmethemoglobin standard (equivalent standard hemoglobin concentration is 15.1 g/dl) with Drabkin’s solution. While validating the microfluidic platform, 2 μl of blood was mixed with Drabkin’s solution (23RR621-80) in a volumetric ratio of 1:250, to yield cyanmethemoglobin in micro-centrifuge tubes, before being subsequently added to different hydrophilic sites of the microfluidic platform, pre-wetted with 2 μl of chromogenic reagent. The entire methodology of detection has been summarized in Figure 1. The extent of colorimetric reaction at the “reaction pads” was monitored by acquiring images at an interval of one second by a digital camera (Nikon D5200).

### Analysis of data

The images of the hydrophilic-reaction sites were processed using the open online program ImageJ (https://imagej.nih.gov/ij/). We have chosen a time window of 150-210 seconds from the reaction kinetics for selecting and processing images for our analysis. A linear transformation was judiciously employed to account for fluctuations in magnification, focusing and illumination, resulting in mapping the intensities of black (*I*_*BR*_ & *I*_*BUR*_) and white (*I*_*WR*_ & *I*_*WUR*_) regions of the microfluidic device to 0 and 255 respectively. The subscripts ‘R’ and ‘UR’ denote the intensities corresponding to the reacted and unreacted reaction pads respectively. The intensity of the image of the reaction pad, wetted with the chromogenic reagent, before the onset of chemical reaction (*I*_*UR*_) was first calculated and linearly transformed. Dynamic variation of intensity of the reaction pad during the course of the chemical reaction was explicated and discussed in detail in the "Results and Discussion". The temporal average of the intensity (*I*_*R,MAX*_) in an appropriate timeframe, after the completion of the chemical reaction, was subsequently obtained and scaled accordingly. The difference of the scaled intensities (Δ*I*), corresponding to a given concentration, was obtained by: 
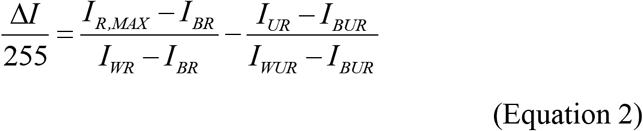

### Statistical Analysis

All statistical analyses were performed using MATLAB 2017. Pearson correlations, linear regression models, and prediction bounds were employed to estimate the correlation between Hgb levels via pathological hematology analyzer and POC device. Sensitivity and specificity results were computed for the entire set of patients, irrespective of the age group.

### Study approval

An approval of ethical clearance was taken from Institute Ethical Committee (IEC) for the commencement of this study. Pediatric and adult patient blood samples were obtained from the pathology clinic of the B.C. Roy Technology Hospital. All specimens were collected after informed approval was received from patients or their parents/guardians. Each consented patient only provided one sample on the day of consent.

## Author Contributions

SC conceptualized and envisioned this study. SKB and SB were responsible for the design, fabrication, experiments, post-processing, and analysis. SK was involved in technical design and strategy for execution of the study with SKB and SB. NKS actively involved in the clinical validation part of the work. All members were involved in technical discussion in different stages of the project and contributed to manuscript writing.

## Acknowledgements

This work is a part of the project entitled “Development of Smartphone Integrated Generic Microfluidic Devices for Rapid, Portable and Affordable Point-of-Care Diagnostics” (Project ID: 4429) (GDD), funded by a grant from the Ministry of Human Resource Development (MHRD) and Indian Council of Medical Research (ICMR), Department of Health Research, Ministry of Health and Family Welfare, New Delhi. This material is based upon work supported by the MHRD Research Fellowship from Government of India, MHRD, Department of Higher Education, New Delhi. We appreciate the collaborative support from BC Roy Technology Hospital, Indian Institute of Technology for providing blood samples and their pathological test results.

